# Sub-stoichiometric Hsp104 regulates the genesis and persistence of self-replicable amyloid seeds of a yeast prion protein

**DOI:** 10.1101/2021.03.08.434509

**Authors:** Sayanta Mahapatra, Anusha Sarbahi, Priyanka Madhu, Hema M. Swasthi, Samrat Mukhopadhyay

**Affiliations:** Centre for Protein Science, Design and Engineering, Indian Institute of Science Education and Research (IISER) Mohali, Punjab, India; Department of Biological Sciences, Indian Institute of Science Education and Research (IISER) Mohali, Punjab, India; Department of Chemical Sciences, Indian Institute of Science Education and Research (IISER) Mohali, Punjab, India

## Abstract

The prion-like self-perpetuating conformational conversion is involved in both transmissible neurodegenerative diseases and non-Mendelian inheritance traits. The transmissibility of amyloid-like aggregates is dependent on the stoichiometry of chaperones such as heat shock proteins. To provide the mechanistic underpinning of the generation and persistence of prefibrillar amyloid seeds that are critical for the prion-like propagation, we studied the effect of Hsp104 disaggregase on the assembly mechanism of a yeast prion determinant of *Saccharomyces cerevisiae* Sup35. At low sub-stoichiometric concentrations, Hsp104 exhibits a dual role and considerably accelerates the formation of seeding-competent prefibrillar amyloids by shortening the lag phase but also prolongs their persistence by introducing unusual kinetic halts and delaying their conversion into matured fibers. Hsp104-mediated amyloid species comprise a more ordered packing and display an enhanced autocatalytic self-templating ability compare to amyloids formed without Hsp104. Our findings underscore the key functional and pathological roles of sub-stoichiometric chaperones in prion-like propagation.

## Introduction

Protein misfolding results in the deposition of proteinaceous β-rich amyloid aggregates and is associated with a range of fatal neurodegenerative diseases.^1,2^ Prions belong to one of the sub-classes of amyloids that can exhibit a self-perpetuating conformational conversion. They can migrate from a small infected patch to the distal parts of the neuronal tissues resulting in adverse cellular consequences leading to neurodegeneration.^3^ The prion-like mechanism has also been proposed for other amyloidogenic proteins such as α-synuclein, tau, amyloid β (Aβ), Huntingtin, p53 and so forth.^4,5,6,7,8,9^ The prion-like spreading in the brain is thought to involve the preformed self-replicable amyloid entities called the propagons or seeds that are considered as the minimum units of amyloid infections. Aging increases the frequency of these events as the protein quality control (PQC) system faces challenges.^10^ The PQC system comprising a sophisticated network of proteins, called the chaperones, is not only devoted to the proper folding of nascent polypeptide chains but also guides the unfolded and misfolded proteins to attain the native three-dimensional shape by inhibiting their aberrant aggregation and eliminating irreversibly aggregated proteins.^11,12,13^ Disaggregases (Hsp110 in higher eukaryotes; ClpB in *E .coli*; Hsp104 in yeasts) belong to an important class of chaperones that are involved in the ATP-dependent and cochaperone-regulated disassembly of aggregated proteins that bypass the other surveillance of the PQC system.^14,15,16,17^ In aged neurons, however, one of the critical manifestations of the PQC dysfunction is the lower expression of these disaggregases such as Hsp110, Hsp70, and so forth. The insufficiency of the disaggregases has been linked with the amyloid-promoting propensity and its fatal consequences.^18^

A functional prion protein (Sup35) that is beneficial to yeast serves as an excellent model to develop the prion concept as well as to elucidate the role of disaggregases in the prion-like transmission of several disease-associated amyloids. This is due to the following reasons.^19^ Firstly, due to its super-structural resemblance with disease-linked amyloids that exhibit a prion-like propagation. Secondly, the distinct prion phenotypes [*PSI*^+^ and *psi*^-^] and the prion strains can be recapitulated by the protein-only transmission using *in vitro* generated amyloids. Interestingly, these traits exhibit a dose-dependence with respect to the cellular disaggregase machinery of yeasts namely, Hsp104, a hexameric AAA+ ATPase that controls the cross-generational non-Mendelian inheritance of [*PSI*^+^] phenotype and is reminiscent of the neuron-to-neuron transmission of self-replicable amyloid seeds.^20,21,22^ At higher concentrations, Hsp104 dissolves the aggregates up to the non-infectious level and impair the passage of the prion phenotype resulting in curing of the [*PSI*^+^] phenotype.^23^ Also, the genetic or chemical inactivation of Hsp104 hinders the propagation of [*PSI*^+^] phenotype due to the unavailability of enough prefibrillar seeds that are generated from matured fibrils by Hsp104.^24,25^ The generation and persistence of prefibrillar amyloids as the seeds for the successful prion-like infection is critical as matured amyloid fibrils show limited infective potential due to their fewer ends of polymerization and lower cytoplasmic diffusibility.^26,27,28^ Understanding the underlying mechanism of prion formation and propagation at low concentrations of Hsp104 is important for the studies mimicking aged neurons under-expressing disaggregases and concomitantly exhibiting the prion-like amyloid colonization. The studies have suggested that the sub-stoichiometric Hsp104 accelerates the fibrillation that minimizes the persistence of prefibrillar aggregates and forms inefficient fibrillar seeds.^29^ Also, the chance of curetting prefibrillar seeds indirectly through disaggregating fibrils by Hsp104 in such low concentrations is very nominal that collectively fails to explain the critical aspect of the abundance of prefibrillar amyloid species that is crucial in the self-templating cascade of prions.^30^

In this work, to uncover the molecular mechanism behind the feasible generation of prefibrillar amyloids as effective seeds by the low concentrations of Hsp104 through the *in vitro* recapitulation of the cellular scenario of yeasts in a minimalistic approach, we used the NM domain of the *Saccharomyces cerevisiae* translation termination factor, Sup35. The NM domain is intrinsically disordered in the (monomeric) non-prion form and comprises the N-terminal part abundant in polar uncharged amino acids (glutamine, asparagine, and tyrosine) and a highly charged middle region (M) (Figure 1a). The NM domain of Sup35 is necessary and sufficient to recapitulate all the characteristics of the prion state, and therefore, represents a prion determinant in yeast. Using sub-stoichiometric ratios of Hsp104, we detected a pronounced kinetic alteration of the NM aggregation behavior that supported not only the rapid generation of seeding competent prefibrillar amyloids but also ensured the prolonged persistence of these species before their recruitment into matured amyloid fibers. Additionally, we were also able to capture conformationally distinct, Hsp104-remodeled NM species that exhibit a much higher seeding potential.

**Figure 1.**
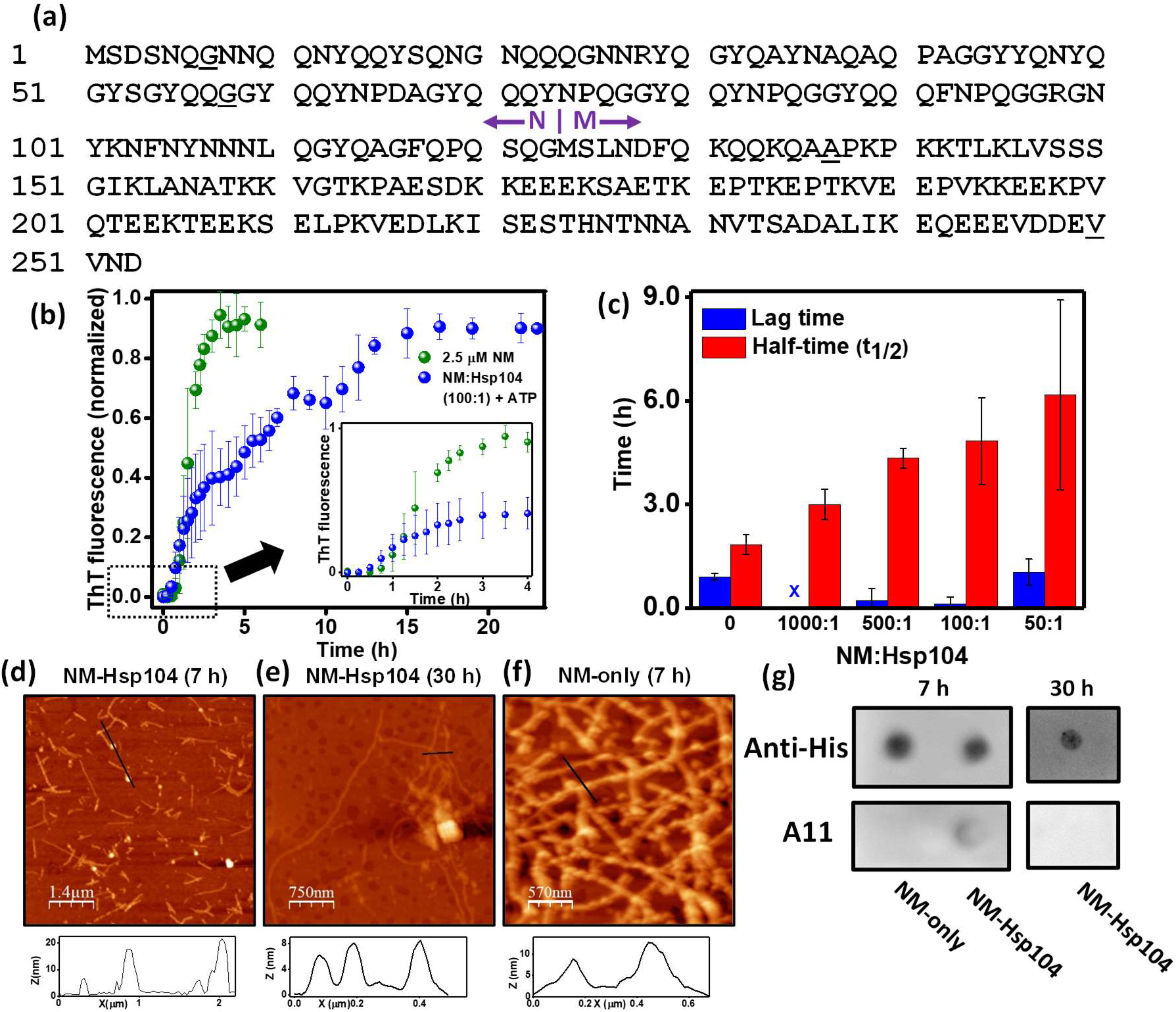
(a) The amino acid sequence of Sup35NM showing the putative boundary between the N- and M-domain. Residue positions for single Trp variants are shown as underscored. (b) Normalized ThT fluorescence kinetics of NM (2.5 μM) without or with Hsp104 (0.025 μM) and ATP (5 mM) during amyloid formation (stirred at 80 rpm at room temperature). The kinetics of the first five hours from the commencement of the reactions are shown in the inset. The individual ThT kinetics of NM-Hsp104 aggregation is shown in figure S1a-c. (c) The lag time and t_½_ of the NM aggregations without or with various sub-stoichiometric ratios of Hsp104. The lag times were retrieved by fitting the first 6 h fluorescence intensities to sigmoidal function and t_½_ were determined from the time-points when the normalized fluorescence intensities reached 0.5. The standard errors were estimated from three independent experiments. (d,e) AFM images of NM amyloids (2.5 μM monomers) showing the oligomers and protofibrils with height of ~ 20 nm and 7 nm, respectively, in the presence of Hsp104 (0.025 μM), plus ATP after 7 h (d) and fibrils with the height ~ 7 nm after 30 h (e) and also after 2 h and 25 h (Figure S1g,h) from the commencement of the reactions. (f) AFM image of NM fibrils (2.5 μM monomers) formed after 7 h of aggregation in the absence of Hsp104 with the height ~ 9 nm. (g) Samples from the NM aggregation reactions without or with Hsp104 (NM: Hsp104 100:1) and ATP were spotted on nitrocellulose membrane after 7 h from the commencement of the reactions and dot-blotted with the anti-His and A11 antibodies. The sample was also spotted from the Hsp104-mediated NM aggregation reaction after 30 h of polymerization.

## Results

### Hsp104 modulates the NM assembly kinetics

We first carried out the amyloid formation kinetics at a low micromolar protein concentration in the absence of Hsp104 using a well-known amyloid reporter namely, thioflavin-T (ThT).The aggregation of NM (2.5 μM) proceeded via typical nucleation-dependent polymerization kinetics possessing a lag phase of approximately 50 min, an assembly phase, and a saturation phase (Figure 1b).^31,32^ In order to investigate the effects of the low concentrations of Hsp104 in NM assembly, we performed the aggregation kinetics in the presence of Hsp104 at several sub-stoichiometric ratios containing ATP and an ATP regeneration system. We observed rapid oligomerization of NM and shortening of the lag phase in the presence of Hsp104, an observation that is consistent with the previous study. ^29^ At the lowest concentration of Hsp104 (NM:Hsp104 = 1000:1), the lag phase is almost abolished (Figure 1b,c and S1e). Interestingly, shortening of the lag phase in the presence of a low concentration of Hsp104 is associated with a delay in the assembly phase. This observation indicated that the assembly and maturation of Hsp104-induced early prefibrillar species get retarded in a dose-dependent manner (Figure 1c and S1f). As a control experiment, we performed NM aggregation just in the presence of ATP and ATP regeneration system and observed no significant change in the NM aggregation profile in the absence of Hsp104 (Figure S1d). Next, in order to directly visualize the nanoscale morphology, we carried out atomic force microscopy (AFM) imaging, which revealed the appearance of a mixture of spherical oligomers and protofibrils in the presence of Hsp104. In contrast, in the absence of Hsp104, we observed primarily matured fibrils at the same timepoint (7h) (Figure 1d-f). To further reconfirm the spherical entities as amyloid oligomers, we probed the NM-Hsp104 aggregation reactions using A11 antibody that is specific for oligomers.^33,34,35^ In the sample spotted after 7h from the commencement of the reaction, we detected a weak signal indicating the existence of a low fraction of oligomeric species in the presence of a low concentration of Hsp104 but not in the absence of Hsp104. This weak A11 activity in the presence of Hsp104 disappeared after 30h due to the conversion of all oligomeric species into matured fibrils (Figure 1g). Together, this set of results showed Hsp104, at a sub-stoichiometric ratio, accelerates the early oligomerization events resulting in the formation of prefibrillar species but decelerates the growth kinetics allowing a prolonged persistence of prefibrillar species before they mature into amyloid fibers.

### The role of Hsp104-mediated disaggregation in the NM assembly kinetics

Next, we asked whether the modulation in the aggregation kinetics by Hsp104 is due to its specific disaggregase activity or a passive perturbation in the NM polymerization by this chaperone. In order to distinguish between these two possibilities, the aggregation reaction of NM monomers with Hsp104 was set up in the absence of ATP, as Hsp104 is ATP-dependent amyloid disassembling machinery, and we did not observe any measurable change in the aggregation kinetics (Figure 2a). We also performed NM aggregation with Hsp104 and ATP, but in the presence of a millimolar concentration of GdmCl that acts as a potent inhibitor of Hsp104 by preventing its ATP hydrolysis-dependent disaggregase activity.^36^ In this case also, we did not observe any changes in the aggregation profile that suggested a coordinated role of ATPase- and disaggregase activities of Hsp104 in the alteration of the NM aggregation behavior (Figure 2b). On this basis, we further tested if there was a different extent of monomer recruitment in amyloids in NM-only and Hsp104-mediated NM assembly due to their pronounced kinetic dissimilarities. However, when we retrieved the high molecular weight aggregates in the pellet fraction after the completion of the aggregation reactions by high-speed centrifugation and monomerized them using the denaturant, we observed similar intensities of the NM band on SDS-PAGE for NM-only and Hsp104-mediated NM aggregations. This indicates the recruitment of nearly the same fraction of NM monomers into aggregates in both types of reactions (Figure 2c). By the scrutiny of the aggregation profiles, we noticed some temporary halts resulting in separable biphasic kinetics (marked in Figure 2b) in the amyloid formation in NM-Hsp104 aggregation reactions, more pronounced in the relatively higher ratios of Hsp104, and often these halts in the aggregation are reported to be associated with the fresh recruitment of monomers to form more amyloids on the surfaces of already existed aggregates by the mechanism called ‘secondary nucleation’(Figure S1f).^37^ To assess the possibility of secondary nucleation here, we aliquoted the NM-Hsp104 reaction mixture just before the halt and after the completion of the aggregation, and then the retrieved amyloids were monomerized using the denaturant. The nearly identical monomeric NM band intensities in the SDS-PAGE in both pre-halt and post-halt samples ensured no additional recruitment of monomers via secondary nucleation (Figure 2d). All-inclusive, this set of data suggested the specific ATP-dependent GdmCl-sensitive enzymatic activity of Hsp104 in the modulation of NM assembly kinetics resulted from the tussle between the intrinsic propensity of the NM amyloids to mature into higher-order aggregates and Hsp104 mediated amyloid disaggregation.

**Figure 2.**
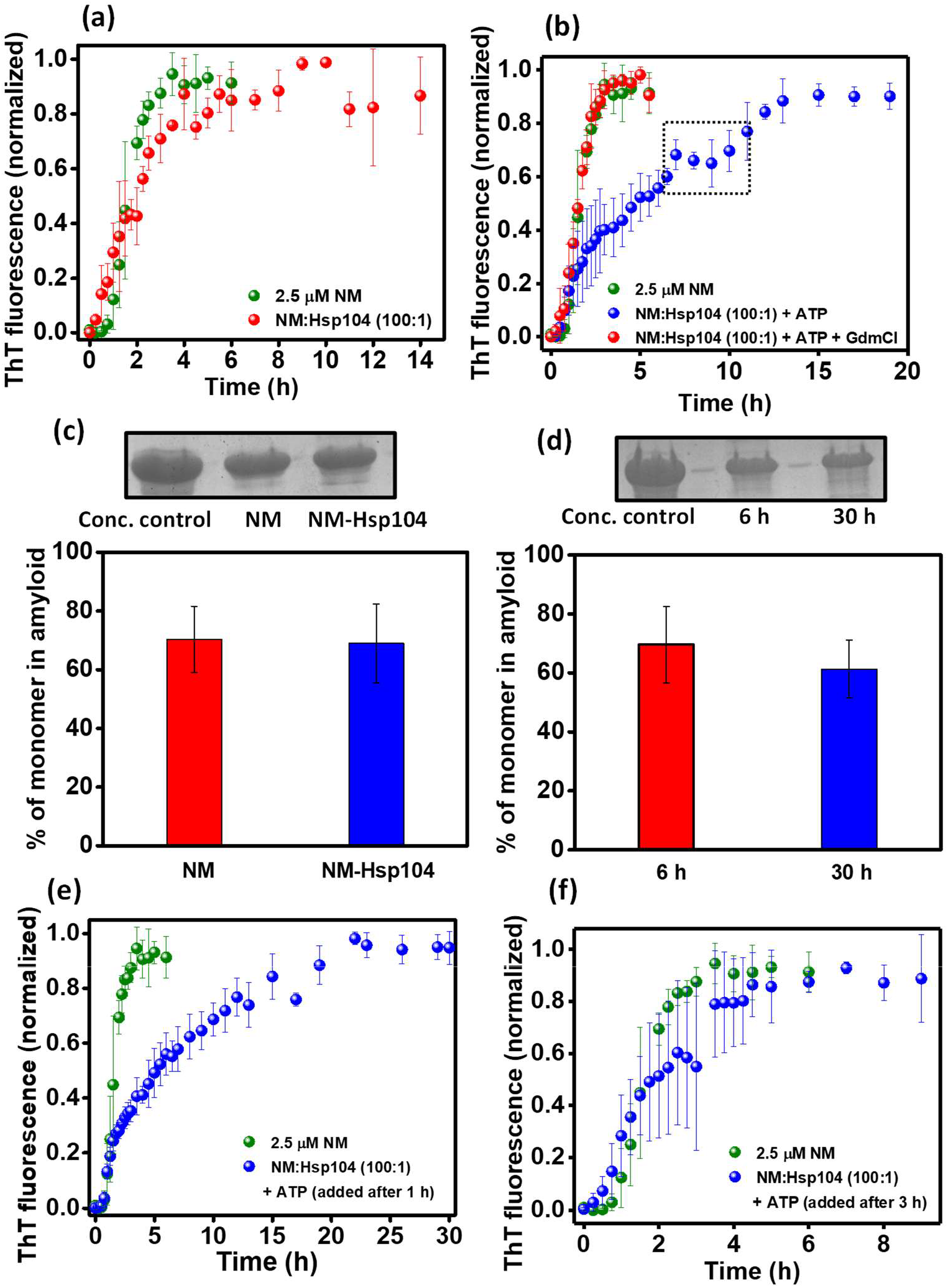
(a) Normalized ThT fluorescence kinetics of rotated (80 rpm) NM (2.5 μM) aggregation without or with Hsp104 (0.025 μM). (b) Normalized ThT fluorescence kinetics of rotated (80 rpm) NM (2.5 μM) aggregation without or with Hsp104 (0.025 μM), plus ATP and with Hsp104 (0.025 μM) and ATP, in the presence of GdmCl (3 mM) in the assembly buffer that does not alter the NM aggregation (Figure S2a). Representative ‘halt’ in the NM-Hsp104 aggregation is marked. (c) The relative quantification by ImageJ software of NM monomers retrieved from the amyloids formed from the rotated (80 rpm) polymerization of NM (2.5 μM) without or with Hsp104 (0.025 μM) and ATP for 6 h or 30 h, respectively, with respect to the concentration control in SDS-PAGE. (d) The relative quantification by ImageJ software of NM monomers retrieved from the amyloids in the aliquots of rotated (80 rpm) polymerization of NM (2.5 μM) with Hsp104 (0.025 μM) and ATP, aliquoted before the ‘halt’ marked in Figure 2b (6 h from the commencement of the aggregation) and at the end of the polymerization (30 h from the commencement of the aggregation). (e,f) Normalized ThT fluorescence kinetics of rotated (80 rpm) NM (2.5 μM) aggregation without or with Hsp104 (0.025 μM), plus ATP, introduced after (e) 1 h (f) 3 h from the commencement of the reaction. (All the standard deviations were calculated from three independent experiments.)

### The role of Hsp104 in the amyloid maturation

During the budding process of yeasts, the daughter cells receive a fraction of the cytoplasm from their mothers containing the preformed amyloid species of various molecular weights. Inefficient conversion of these low molecular weight aggregates into matured fibers is critical to maintain and propagate the amyloid-linked [*PSI*+] phenotype because of the limited infectivity of the mature fibers. Therefore, in order to delineate the putative role of Hsp104 in the persistence of low molecular weight amyloid species, we introduced a low concentration of Hsp104 with ATP at the late-lag phase (0.5 h after the commencement of the reaction) and at the early log phase (1 h after the commencement of the reaction) of NM aggregation reactions (Figure 2e and S2b,c). The kinetics revelated that Hsp104 delayed the maturation of these already formed particles to the higher-order amyloids. In contrast, when Hsp104 and ATP were introduced at the end of the elongation phase (3 h after the commencement of the reaction) of the NM aggregation reactions, no significant modulation in the aggregation kinetics was observed. This observation revealed the inability of the low concentration of Hsp104 to manipulate the higher molecular weight aggregates (Figure 2f). Together, these results led us to surmise that the low concentration of Hsp104 enhanced the abundance of low molecular weight species as opposed to mature fibrils not only by amending the *de novo* aggregation but also decelerating the conversion of the preformed amyloid species to higher molecular weight fibers.

### The seeding capability of the Hsp104 remodeled amyloid

Not only the abundance but also the prefibrillar particles need to effectively seed the amyloid polymerization to continue the cycle of typical prion-like amplification for the inheritance of the [*PSI*^+^] phenotype. To shed light on this critical aspect of prion inheritance, we studied the seeding capability of the amyloid species of Hsp104-mediated NM aggregation by aliquoting preformed amyloids from the reaction mixture at different time points (Figure 3a). The NM-Hsp104 aggregation mixture was introduced into the fresh NM-only polymerization reactions in the buffer containing GdmCl to suppress the effect of Hsp104. This allowed us to study the effect of Hsp104-remodeled aggregates in seeding the NM-only aggregation kinetics without the influence of Hsp104 in seeded reactions. This set of studies showed that the amyloid prefibrillar entities of Hsp104-induced NM-aggregation had a greater potential to accelerate the fresh NM fibrilization compare to the amyloids of NM-only aggregation reactions as reflected in the lag time and the half-time (t_½_) of the seeded kinetics (Figure 3b-d and S3a-c). Moreover, the seeding ability of the particles of NM-Hsp104 aggregation reactions that aliquoted after 25 min demonstrated the early appearance of the seeding-competent amyloid species in Hsp104-mediated NM aggregation compared to the NM-only aggregation reaction. This observation indicated that the source of this early seeding ability lies in the composition of the amyloid particles of the NM-Hsp104 aggregation, which was enriched in the precursor of fibrillar amyloids of NM having a better seeding potential as opposed to matured fibrils. We also surmise that the intermediate NM particles demonstrated higher seeding potential than matured fibers (Figure 3e). However, intriguingly, even the NM-Hsp104 fibrils also displayed better seeding potential compare to the NM fibrils (Figure 3f). Taken together, the greater capability of Hsp104-designed NM fibrils to catalyze the NM assembly than the typical NM fibrils established the fact that the reason behind their better seeding ability was not only related to the polymerization hierarchies and sizes of the amyloids present in the seed-aliquots but also the conformational attribute of Hsp104-remodeled fibrils. Therefore, we postulated that Hsp104, at sub-stoichiometric concentrations, can craft structurally altered amyloids that can autocatalytically accelerate a fresh aggregation reaction more efficiently owing to their distinct amyloid packing. Next, we aimed at distinguishing the conformational characteristics of NM-only and NM-Hsp104 amyloids by following an array of distinct biochemical and biophysical readouts.

**Figure 3.**
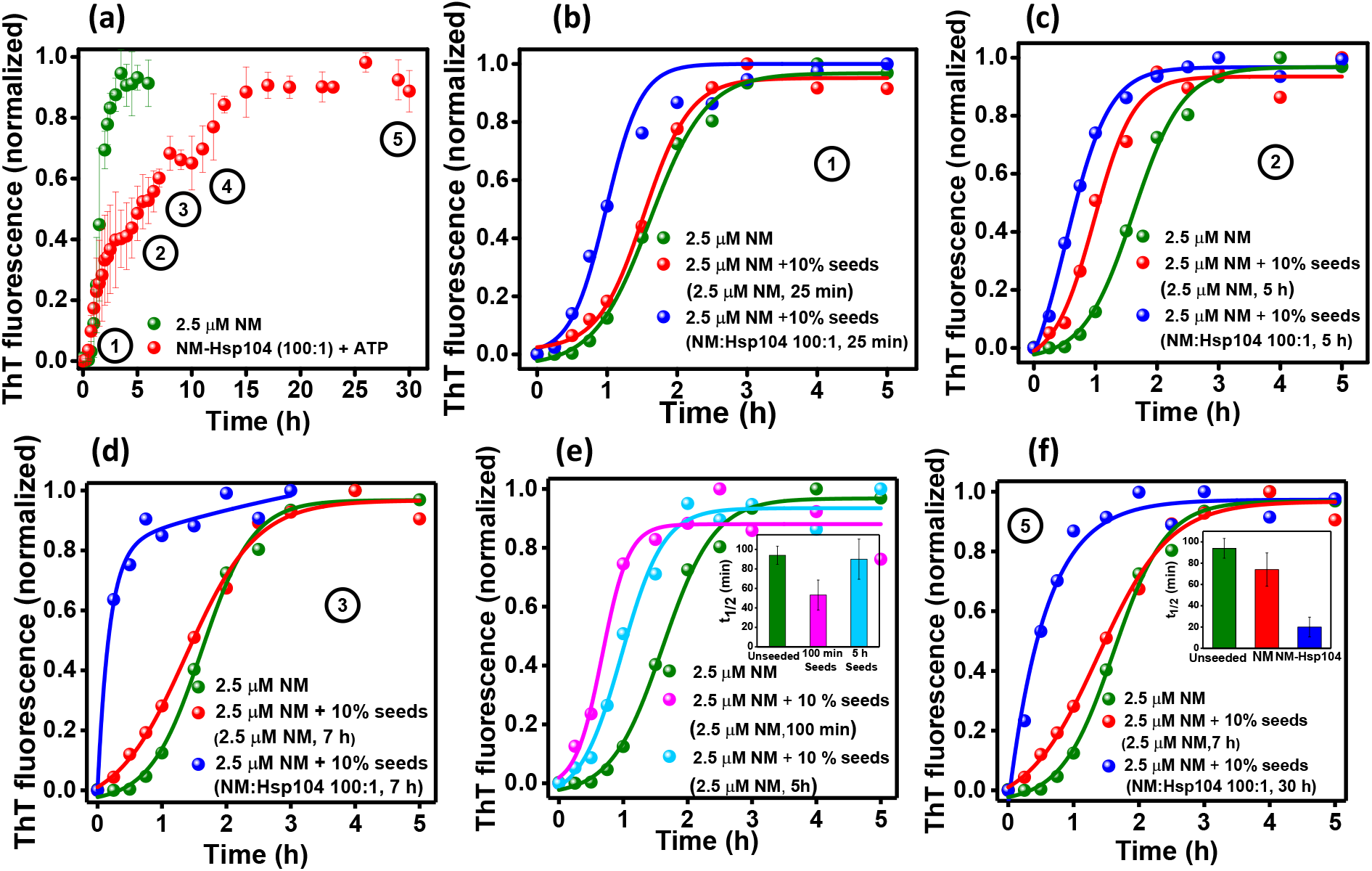
(a) Normalized ThT fluorescence kinetics of rotated (80 rpm) NM (2.5 μM) aggregation without or with Hsp104 (0.025 μM), plus ATP, and aliquots were withdrawn from these reactions as seeds at the indicated time-points (circled numbers) and introduced to the fresh aggregation of NM (2.5 μM) in assembly buffer containing GdmCl (3 mM). (b-d) Representative normalized ThT fluorescence kinetics of rotated (80 rpm) NM (2.5 μM) aggregation without or with 10% (w/w) seeds of NM-Hsp104 or NM aggregation which were aliquoted after (b) 25 min (c) 5 h (d) 7 h, and after 10 h (Figure S3a) from the commencement of the aggregation reactions. (e) Representative normalized ThT fluorescence kinetics of rotated (80 rpm) NM (2.5 μM) without or with 10% (w/w) seeds from NM aggregation that were aliquoted after 100 min and 5 h showing the t_½_ of the unseeded and seeded aggregations. Standard deviations were calculated from three individual experiments (inset). (f) Representative normalized ThT fluorescence kinetics of rotated (80 rpm) NM (2.5 μM) aggregation without or with 10% (w/w) seeds from NM or NM-Hsp104 aggregation that were aliquoted after 7 h or 30 h, respectively. t_½_ of the unseeded and seeded aggregation reactions are shown in the inset. Standard deviations were calculated from three individual experiments (inset).

### Hsp104 induces amyloid structural diversity

In order to monitor the amyloid structural diversity, we first studied the SDS-solubility of NM-only and NM-Hsp104 aggregates. The SDS-induced thermal denaturation was earlier used to identify the structural diversity by monitoring the dissimilar thermal stability of two different yeast prion strains generated *in vitro*^38^ We fibrilized NM without or with Hsp104 at the sub-stoichiometric ratio and then treated these fibrils with 2% SDS and heated from 25 °C to 100 °C.

Then we quantified the monomeric fraction derived from this treatment on SDS-PAGE (Figure 4). The temperature-dependence of the monomeric fraction exhibited a sigmoidal profile showing an increase in the monomeric population with increasing temperature in the two types of fibrils. The dissimilar melting temperatures (T_m_) of NM-only and NM-Hsp104 fibrils revealed altered thermal stability arising due to their distinct supramolecular structural differences despite having similar nanoscale morphologies. However, NM-Hsp104 fibrils prepared in the presence of GdmCl that acts as an inhibitor of Hsp104 exhibited thermal stability that is similar to NM-only fibrils. These results together indicated that the disaggregation-competent Hsp104 induces an altered amyloid packing of NM compared to pure NM fibrils.

**Figure 4.**
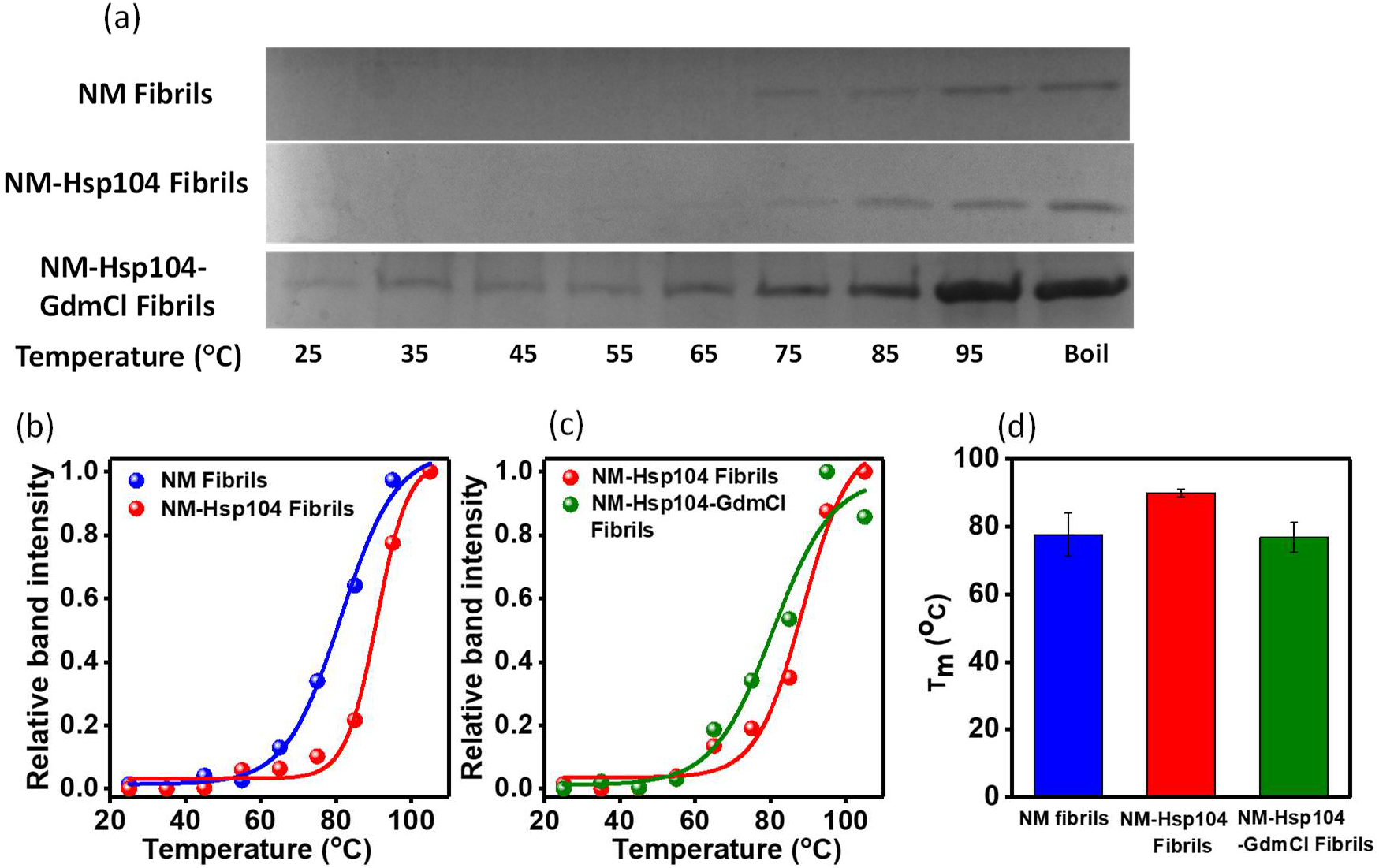
(a) SDS-PAGE gel image of concentrated fibrils formed from the NM monomers (2.5 μM) that fibrilized in the absence or presence of Hsp104 (0.025 μM) and ATP and also in the presence of Hsp104 (0.025 μM), ATP and GdmCl (3 mM) which were then heated with SDS-PAGE loading dye (2% SDS) for 5 min at indicated temperatures. (b,c) The fitted band intensity of the monomers melted from (b) NM and NM-Hsp104 fibrils, (c) NM-Hsp104, and NM-Hsp104-GdmCl fibrils with temperature to sigmoidal functions after relatively quantified using ImageJ software. (d) Melting temperature (T_m_) of NM, NM-Hsp104, and NM-Hsp104-Gdmcl fibrils retrieved from the midpoint of the sigmoidal melting curve. Standard deviations were calculated from three individual experimental replicates.

### Hsp104 alters fibril fragility and protease digestion profiles

To further support our assertion that Hsp104 induces conformationally altered fibrillar architecture, we intended to distinguish NM-only and Hsp104-mediated fibrils by their fragility. We fragmented the NM-only and Hsp104-designed NM fibrils using three different ways: Fragmenting fibrils by ultrasonic sound, by incubating fibrils at a high concentration of Hsp104, and by keeping the fibrils under unagitated conditions at room temperature for 24 h.

Irrespective of the fragmentation method, the ThT fluorescence exhibited dissimilar kinetics for NM-only and NM-Hsp104 fibrils indicating their altered structural packing in these two types of fibrils because the distinct supramolecular arrangements within the fibrils can give rise to the observed kinetic difference in the fragmentation propensity (Figure 5a,b and S4a). Next, in order to further probe into the kinetic stability of these two types of NM fibrils, we performed protease digestion assays. Despite having a generic cross β-sheet secondary structure in different amyloid variants of a given protein, the varied supramolecular packing and nanoscale organization of the monomeric polypeptide units in the polymeric architecture lead to altered sensitivity to proteolytic digestion.^39^ We incubated NM-only and NM-Hsp104 fibrils with an increasing concentration of proteinase K and observed a different digestion pattern on SDS-PAGE and by western blot analysis. In the case of NM-only fibrils, an intact monomeric NM band was visible only in the presence of the lowest amount of protease, as the higher ratios of proteinase K completely digested NM into low molecular weight peptides. In contrast, the NM fibrils designed by Hsp104 showed much more resistance towards protease digestion, and the appearance of the undigested monomeric NM bands at relatively higher concentrations of proteinase K suggested the existence of a more kinetically stable amyloid core in NM-Hsp104 fibrils compared to NM-only fibrils (Figure 5c and S4b).

**Figure 5.**
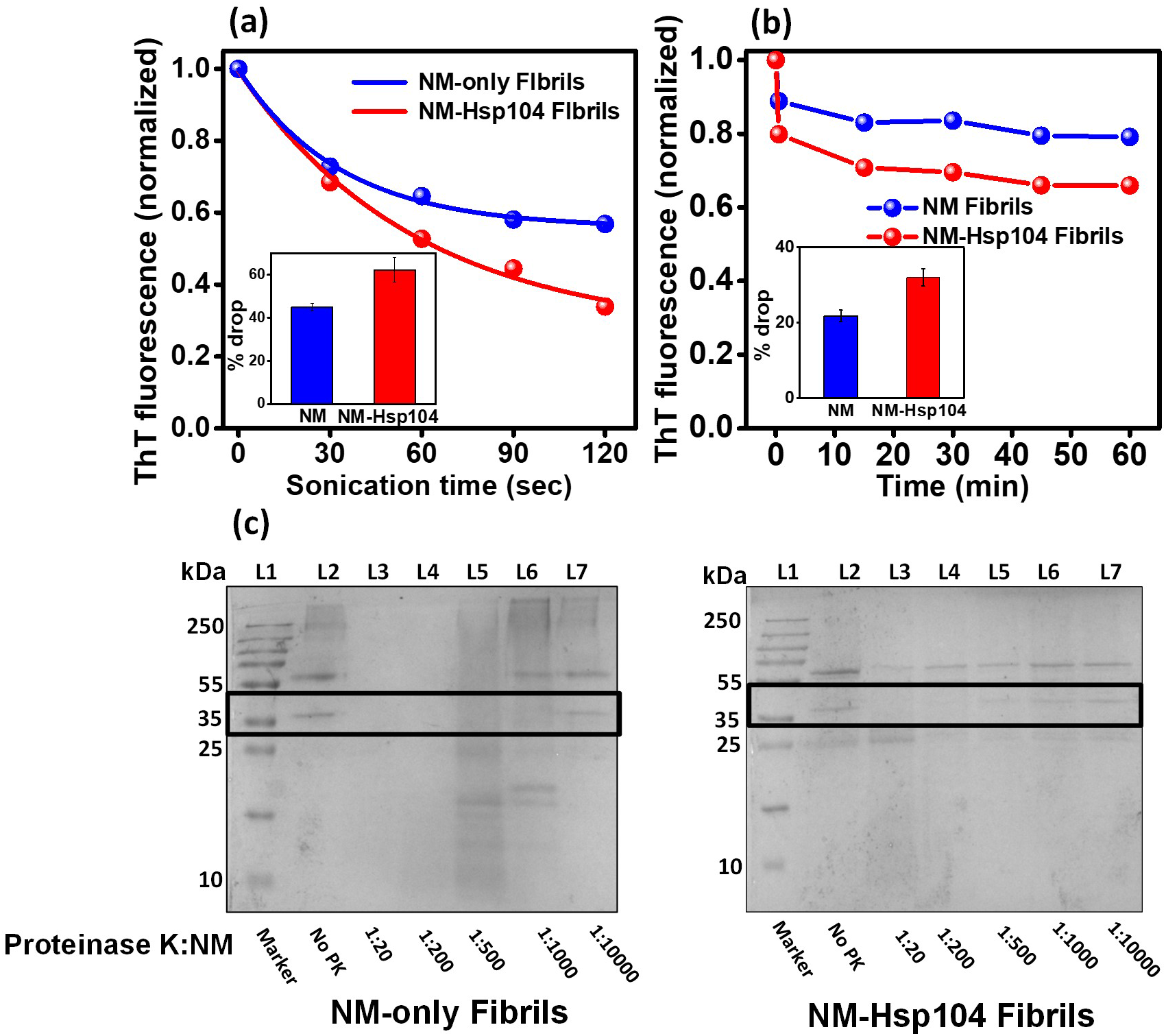
(a,b) Monomeric NM (2.5 μM) was fibrilized without or with Hsp104 (0.025 μM), plus ATP, and then disaggregated by two methods. (a) Representative disaggregation kinetics using the ultrasonic sound pulse of amplitude 5 for several pulses of 30 seconds and the drop in the ThT fluorescence was recorded after each pulse. The ThT fluorescence intensities were normalized with respect to the initial ThT fluorescence intensity. The extent of disaggregation at 80 rpm was estimated from three experimental replicates to calculate the standard deviation (inset). (b) Representative disaggregation kinetics by Hsp104 (0.5 μM), ATP (5 mM), and drop in ThT fluorescence with time at 80 rpm were recorded. The ThT fluorescence intensities were normalized and the extent of disaggregation was estimated as mentioned above (inset). (c) The concentrated fibrils formed from monomeric NM (2.5 μM) without or with Hsp104 (0.025 μM), plus ATP were incubated at 37 °C for 30 min with multiple concentrations of proteinase K followed by the SDS-PAGE analysis. The bands corresponding to the NM monomers are marked with black boxes. Pyruvate kinase was added after the fibrillation in the case of NM-only aggregation to make the reaction mixtures comparable to the NM-Hsp104 aggregation reaction.

### Site-specific conformational mobility distinguishes the types of amyloids

Next, in order to directly capture the site-specific structural information within the amyloid conformers, we performed site-specific fluorescence polarization anisotropy measurements that allowed us to monitor the conformational dynamics in two types of NM fibrils. NM and Hsp104 sequences are devoid of any tryptophan (Trp) residue, and therefore, we used single-Trp variants spanning the NM polypeptide sequence (Figure 1a). We then compared the Trp emission spectra in the monomeric and amyloid states and observed a blueshift for all residue positions in the aggregated form with respect to the denatured monomeric form which is in line with our earlier studies (Figure 6a).^40^ In both types of fibrils, NM-only and NM-Hsp104, the extent of the blue shift was more for N-domain residues than for M-domain residues indicating the solvent-excluded environment in N-domain. Moreover, on the comparison between the two different fibrils, we observed a greater extent of blueshift for NM-Hsp104 fibrils. This blueshift is more pronounced in the N-terminal segment containing residue 7 (Figure 6b). This finding indicated that the NM sequence, especially the N-terminal part, experiences more solvent protection in NM-Hsp104 fibrils compared to NM-only fibrils. Next, we performed the steady-state fluorescence anisotropy measurements that report the site-specific rotational flexibility of Trp in the NM sequence.^41,42^

**Figure 6.**
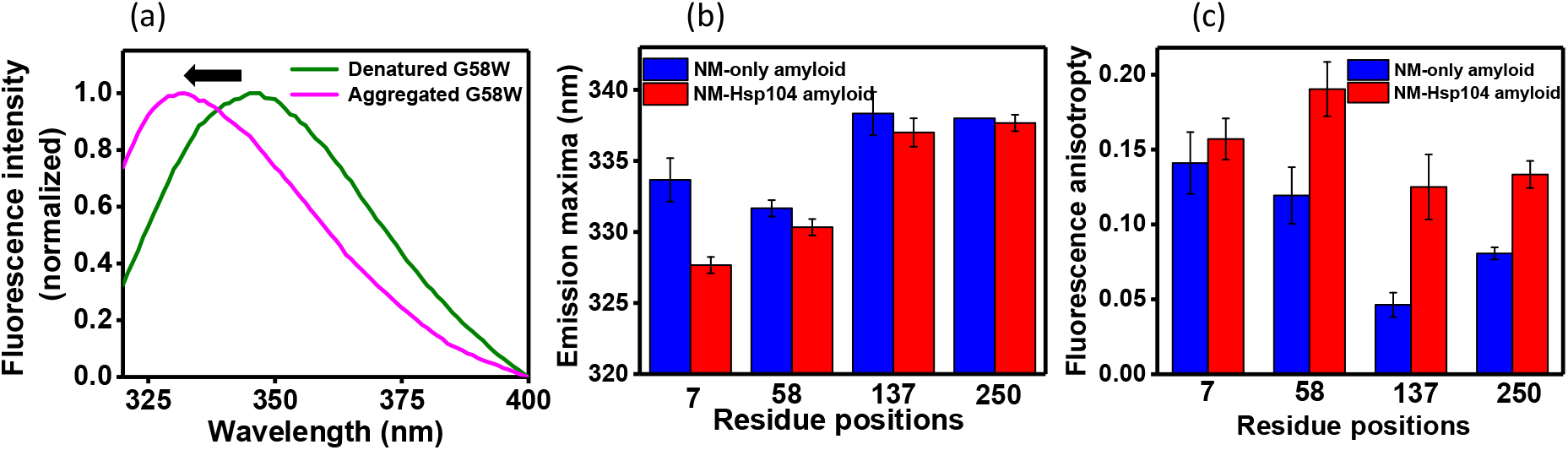
(a) Normalized Trp fluorescence spectra of residue position 58 depicting the blue shift (black arrow) upon conversion into the amyloids. (b) Trp emission maxima of different residue positions in two different amyloid forms, NM, and NM-Hsp104.The excitation and emission slit widths were 1.75 and 6 nm, respectively. (c) Steady-state fluorescence anisotropies of different residue positions in two amyloid states (NM and NM-Hsp104). The standard deviations were estimated from three independent replicates.

Upon conversion to the amyloids, N-domain residues exhibited higher anisotropies indicating the more restricted rotation due to the preferential recruitment of N-domain into the amyloid core. In contrast, the M-segment (residue 137) exhibited a much lower fluorescence anisotropy in both types of amyloids (Figure 6c), an observation that is consistent with previous structural studies on NM amyloid indicating the higher flexibility of the M-segment that is not sequestered into the amyloid core. Additionally, higher fluorescence anisotropy in all residue locations for NM-Hsp104 fibrils compared to NM-only fibrils indicated more closely packed ordered organization in NM-Hsp104 fibrils corroborating our protease digestion results. Taken together, a series of biochemical and biophysical studies supported our hypothesis that apart from the composition of the seeds enriched in prefibrillar aggregates, NM-Hsp104 aggregates, compared to NM-only aggregates comprise an altered and more ordered amyloid packing that allowed them to display their enhanced autocatalytic self-templating ability.

## Discussion

Irrespective of the precise mechanistic differences between the prion-like propagation of neurotoxic species by the transmission of seeds across the cellular membrane and the cross-generational cytoplasmic inheritance of amyloid-associated phenotypic traits by fungal prion particles in the continuous stream of cytoplasm from mother to daughter yeast cells, the successful expedition of infectious amyloid particles into the uninfected cells revolves around the autocatalytic behavior of prefibrillar amyloids.^43,28,26,27^ In this study, by *in vitro* reconstruction, we were able to recapitulate two crucial pro-prion aspects of sub-stoichiometric Hsp104. Firstly, low concentrations of the Hsp104 facilitated the production of seeding-competent amyloid entities and decelerated the conversion of these prefibrillar species into the matured fibrils. Secondly, in addition to the kinetic modulations, we were able to identify conformational remodeling in the amyloids by the Hsp104 contributing to the better seeding potential of these species. Contrary to the view of irreversible hindrance in the fibrillation of various intermediate amyloid species of different amyloidogenic proteins including full-length *S. cerevisiae* Sup35 by Hsp104, here we observed a tradeoff between the two opposing factors namely, the intrinsic nature of NM amyloids to polymerize into higher molecular weight aggregates and the ATP-dependent GdmCl-sensitive disaggregation by Hsp104 that delayed but not inhibited the fibrillation process.^44,45^ Furthermore, the critical balance between these two mutually opposing factors resulted in the observed halts resulting in an apparent biphasic kinetics. These halts allowed a prolonged persistence of the prefibrillar particles, pivotal for the propagation of the prion phenotype.

Apart from ensuring the abundance of highly transmissible, seeding-proficient, fibrillar precursors instead of having fewer and longer matured fibrils with a limited transmissibility, Hsp104, at low sub-stoichiometric concentrations, creates seed units possessing a better seeding potential due to their unique conformational characteristics. Our model in Figure 7 depicts these molecular events demonstrating the rapid formation and slow maturation of lower-order aggregates into the higher-order aggregates in an arbitrarily drawn time-axis in the presence of Hsp104 leading to the abundance and preferential cytoplasmic transmission of conformationally distinct, seeding-proficient oligomers and protofibrils as opposed to the fibrils.^26^ In this study, we probed this conformational identity using a host of biochemical tools that revealed Hsp104-mediated NM fibrils are more stable and contain a more ordered amyloid core compared to NM-only aggregates. In accordance with our biochemical findings, location-specific spectral shifts and dynamics revealed via fluorescence anisotropy measurements indicated more buried locations and higher polypeptide ordering in Hsp104-mediated fibrils. The amyloidogenic N-segment is known to constitute the amyloid core, whereas, the charged M-region possesses some conformational flexibility ^46,40^ as also detected by our fluorescence anisotropy measurements. Hsp104 is known to interact with the M-region of NM, and therefore, can result in the binding-induced restriction in the conformational dynamics of the M-region. However, higher anisotropy in the N-domain in the presence of Hsp104 is likely to be caused by higher structural ordering of the amyloid core in Hsp104-mediated NM fibrils. We would like to note that in contrast to previously reported yeast prion strains, in our case, more stable and ordered NM-Hsp104 fibrils were found to be more fragile to fragmentation by Hsp104 which indicate an intriguing interplay of stability and fragility that can have a diverse phenotypic outcome.^47,48,49,50,51^

**Figure 7.**
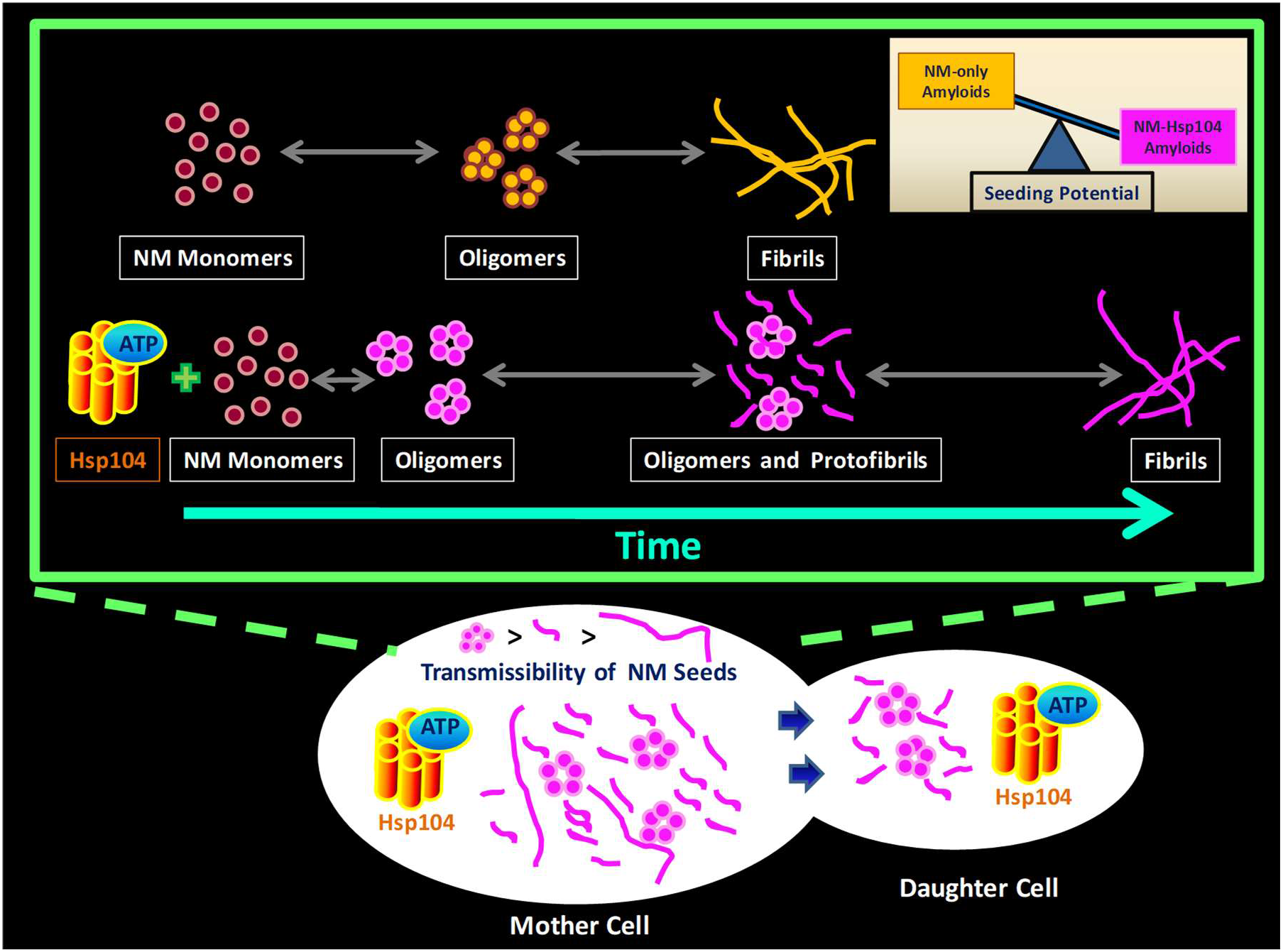
Proposed model for the NM aggregation in the presence of sub-stoichiometric Hsp104. Hsp104 with ATP, as opposed to the long fibrils, ensures the abundance of highly transmissible ^24^ prefibrillar amyloids (oligomers and protofibrils) that shows the greater seeding potential than the amyloids evolved in the absence of Hsp104.

In summary, our findings indicate the pro-[*PSI*^+^] nature of the sub-stoichiometric concentrations of Hsp104, which aid the generation and persistence of the highly transmissible prefibrillar seeding-proficient amyloid species that do not readily transform into matured fibers. The Hsp104-mediated hindrance in the conversion of low molecular weight aggregates into matured fibers is crucial in maintaining and propagating the amyloid-linked [*PSI*^+^] phenotype in yeast. Our results have much broader implications in functional- and pathological prion-like mechanisms in higher organisms. The disaggregase activity and kinetic modulation of Hsps on a wide range of proteins leading to the generation and persistence of conformationally distinct self-replicating amyloid species can potentially underlie a consensus mechanism for the generation and colonization of the toxic, transmissible, neuropathological prefibrillar amyloids. The cascade of molecular events further tuned by the co-chaperones and cellular microenvironments can constitute the prion-like propagation strategies for the distal invasion in the aged brains having impairment in protein homeostasis characterized by the scarcity of disaggregases.

## Materials and Methods

### Materials

HEPES[4-(2-hydroxyethyl)-1-piperazineethanesulfonic acid], Magnesium chloride hexahydrate, Sodium phosphate dibasic dihydrate, tris (hydroxymethyl) aminomethane (Tris), β-Mercaptoethanol, Adenosine-5’-triphosphate disodium salt hydrate (ATP), Dithiothreitol (DTT), Thioflavin-T(ThT) were bought from Sigma (St. Louis, MO, USA). Guanidine hydrochloride (Gdmcl), proteinase-K, and urea were procured from Amresco. Ammonium sulfate, imidazole, lysozyme, sodium dodecyl sulfate (SDS), ethylenediaminetetraacetic acid (EDTA), potassium chloride was bought from HIMEDIA. Potassium hydroxide, sodium chloride, sodium hydroxide, glycerol, A-11 anti-amyloid oligomer antibody, methanol, HRP-conjugated goat-anti-rabbit antibody was procured from Merck. Isopropyl-β-thiogalactopyranoside (IPTG), antibiotics (chloramphenicol and ampicillin) were purchased from Gold Biocom (USA). Enhanced chemiluminescence kit (ECL), HRP-conjugated rabbit-anti-mouse antibody, was obtained from Thermo Fisher Scientific. Ni-NTA column and Q-Sepharose were from GE Healthcare Lifesciences, USA. Phosphoenolpyruvate (PEP) and pyruvate kinase (PK) were procured from Roche Diagnostics, Germany.

### Methods

#### Expression and purification of Sup35NM

C-terminal hexahistidine recombinant Sup35NM proteins were overexpressed in BL21 (DE3) / pLysS cells using IPTG and then from the harvested cells proteins were extracted, the extracted proteins were subjected to first Ni-NTA purification in the gradient of imidazole and further from a Q-sepharose column using the gradient of sodium chloride. The detailed protocol is described by us. ^40^

#### Expression and purification of Hsp104

A modification of a previous protocol was used.^52^ N-terminal His_6_-tag recombinant Hsp104 pPROEX-HTb-Hsp104 of *S. cerevisiae* were overexpressed in BL21(DE3) RIL *E. coli* cells using 1 mM IPTG as inducer at 15 °C for 14 h. Harvested cells suspended in chilled 10 mL lysis buffer (40 mM HEPES-KOH pH 7.4, 500 mM KCl, 20 mM MgCl_2_, 2.5% (w/v) glycerol, 20 mM imidazole) were incubated in 4 °C with lysozyme (2 mg/mL) followed by sonication. The cell debris was removed by centrifugation at 11,500 rpm for 30 min, and the supernatant was subject to Ni-NTA purification using the gradient of imidazole. After Ni-NTA purification, the eluant was buffer exchanged with the (20 mM Tris-HCl pH 8, 0.5 mM EDTA, 5 mM MgCl_2_, 50 mM NaCl) using MWCO 30,000 Amicon Ultra (Millipore) 15 mL centrifugal concentrator units. The protein further purified using the Q-sepharose column using the gradient of NaCl, and the eluant was further buffer exchanged with the cleavage buffer (20 mM HEPES-KOH pH 7.4, 140 mM KCl, and 10 mM MgCl_2_) using the concentrator unit mentioned above. Recombinant TEV protease carrying a hexahistidine tag was used at a ratio His_6_-Hsp104: TEV protease (15:1) to cleave the histidine (His_6_) tag of the Hsp104 at 30 °C for 1 h. The cleaved His_6_ tags of Hsp104, the uncleaved His_6_-Hsp104, and the histidine-tagged TEV protease were removed by binding them with the Ni-NTA resin, and in the flow-through, the pure Hsp104 with no histidine tags were collected and stored in a storage buffer (20 mM HEPES-KOH pH 7.4, 140 mM KCl, and 10 mM MgCl_2_,1 mM DTT, 0.5 mM EDTA) at −80 °C until further use.

#### Amyloid aggregation reaction

For the setting up of aggregation reactions, methanol precipitated NM was dissolved in 8M urea (20 mM Tris-HCl buffer, pH 7.4) for 3 h at room temperature. Monomerized protein was first passed through a 100-kDa filter to remove any pre-existing aggregates if present, and subsequently, the filtrate was concentrated using a 3-kDa filter before the aggregation reaction. The concentrated monomers of NM were further centrifuged at 13,000 rpm for 15 min at room temperature, after which the supernatant was added such that its final concentration is 2.5 μM in assembly buffer (40 mM HEPES-KOH pH 7.4, 150 mM KCl, 20 mM MgCl_2_, 1 mM DTT, 10 μM ThT) without or with Hsp104 and 5 mM ATP and ATP regeneration system (20 mM Phosphoenolpyruvate (PEP), and pyruvate kinase (15 μg/ml) at room temperature under stirring at 80 rpm using the magnetic beads. ThT fluorescence was monitored at room temperature by exciting at 450 nm, and the fluorescence emission was recorded at 480 nm.

Hsp104 influenced NM aggregation reactions were also carried out under the same aggregation conditions and assembly buffer independently with ATP, ATP regeneration system, and 3 mM guanidine hydrochloride (GdmCl) in the assembly buffer and also in the absence of ATP and ATP-regeneration system.

#### Seeded aggregation reaction

Seeds of NM were generated by incubation of monomerized NM (2.5 μM) protein in assembly buffer (40 mM HEPES-KOH pH 7.4, 150 mM KCl, 20 mM MgCl_2_, 1 mM DTT) without or with Hsp104 (0.025 μM), ATP (5 mM) & ATP regeneration system (20 mM PEP, and 15 μg/ml pyruvate kinase) at room temperature under stirring at 80 rpm using magnetic beads. The resulting amyloid seeds from NM or Hsp104 controlled NM aggregation reactions were aliquoted after certain time points from the commencement of the aggregation reactions and added to a 10% (w/w) ratio to the fresh aggregation reaction of NM monomers (2.5 μM) in seeded assembly buffer (40 mM HEPES-KOH pH 7.4, 150 mM KCl, 20 mM MgCl_2_, 1 mM DTT, 10 μM ThT, 3 mM Gdmcl). The seeded aggregation reactions were kept at room temperature under stirring at 80 rpm using magnetic beads, and the ThT fluorescence was recorded with time.

#### Dot-blot assay

Monomeric NM (2.5 μM) was aggregated in the assembly buffer (40 mM HEPES-KOH pH 7.4, 150 mM KCl, 20 mM MgCl_2_, 1 mM DTT) in the presence of Hsp104 (0.025 μM), ATP (5 mM), and ATP regeneration system (20 mM PEP, and 15 μg/ml pyruvate kinase) at room temperature under stirring at 80 rpm, and after 7 h and 30 h from the commencement of the reaction, the aliquots (2 μL) were spotted on the nitrocellulose membrane. NM monomers (2.5 μM) were also aggregated in the absence of Hsp104 and ATP for 7 h under the same conditions in the same assembly buffer and spotted (2 μL) on the nitrocellulose membrane. The blots were blocked using 3% BSA in PBST (0.05 % Tween-20) for 1 h at room temperature and then probed with the primary antibody (A11;1:500) and (Anti-His;1:10,000) overnight at 4 °C. The blots were washed six times with PBST and incubated with appropriate HRP-conjugated secondary antibody for 1 h at room temperature. Again, the blots were washed thrice using PBST and subsequently developed using an ECL kit.

#### Estimation of the Sup35NM monomers recruited in the amyloids

Monomeric NM (2.5 μM) were aggregated in the assembly buffer (40 mM HEPES-KOH pH 7.4, 150 mM KCl, 20 mM MgCl_2_, 1 mM DTT) without or with Hsp104 (0.025 μM), ATP (5 mM), and the ATP regeneration system (20 mM PEP, and 15μg/ml pyruvate kinase) for 6 h or 30 h, respectively, at room temperature under stirring at 80 rpm and the amyloids generated in the reactions were pelleted down at 16,400 rpm for 30 min. The pellets were resuspended in 8 M Urea (20 mM Tris-HCl, pH 7.4) overnight to monomerize the amyloids, and SDS-PAGE was performed. The coomassie stained monomeric NM band intensities were relatively estimated using the ImageJ software concerning the band corresponding to the monomers of a known NM concentration in 8 M Urea (20 mM Tris-HCl, pH 7.4).^53^ To validate the occurrence of secondary nucleation, aliquots were taken from the Hsp104 mediated NM aggregation reaction after 6 h and 30 h, respectively, from the commencement of the aggregation reactions. The amyloids so formed were retrieved and then monomerized in 8 M Urea (20 mM Tris-HCl, pH 7.4) following the protocol mentioned above, and after the SDS-PAGE, the fraction of monomers recruited in the amyloids were compared in both the samples by comparing the coomassie band intensities of the NM monomers using the ImageJ software.

#### Thermal melting of fibrils

Monomeric NM (2.5 μM) was aggregated for 6 h or 30 h in assembly buffer (40 mM HEPES-KOH pH 7.4, 150 mM KCl, 20 mM MgCl_2_, 1 mM DTT) in the absence or presence of (0.025 μM) Hsp104 with ATP (5 mM) and the ATP-regeneration system (20 mM PEP, and 15 μg/ml pyruvate kinase), respectively at room temperature under stirring at 80 rpm using a magnetic bead for the fibril formation. Then, fibrils were passed through a 50-kDa filter to concentrate ~20 times and eliminate the unrecruited monomers. The concentrated fibrils, with SDS-PAGE loading dye (2% SDS), were incubated at different temperatures for 5 min, and then SDS-PAGE was performed. Coomassie-stained bands were quantified using ImageJ software. The band intensities were plotted against the incubation temperatures and fitted to the sigmoidal function. NM (2.5 μM) was also aggregated in the same assembly buffer, additionally having Gdmcl (3 mM) in the presence of the same amount of Hsp104, ATP, and, ATP regeneration system for 6 h for the fibrilization and the thermal melting experiment was performed on these fibrils as well.

#### The proteinase K digestion of fibrils

Monomeric NM (2.5 μM) was aggregated in the assembly buffer (40 mM HEPES-KOH pH 7.4, 150 mM KCl, 20 mM MgCl_2_, 1 mM DTT) in the absence or presence of Hsp104 (0.025 μM), ATP (5 mM), and ATP regeneration system (20 mM PEP, and 15 μg/ml pyruvate kinase) for 6 h or 30 h, respectively, to generate the fibrils and after that, the fibrils were concentrated and freed from unrecruited monomers using a 50-kDa filter. The concentrated fibrils (in the supernatant) from both the aggregation reactions were incubated with proteinase K in multiple ratios at 37 °C for 30 min, and digestion reactions were terminated by adding SDS-PAGE loading dye, and then SDS-PAGE was performed. The undigested NM monomers were also probed with (Anti-His; 1:10,000) antibodies in western blot analysis. Pyruvate kinase (15 μg/ml) was added after fibrillation in the case of NM-only aggregation to make the reaction mixture comparable to the NM-Hsp104 aggregation reaction.

#### Disaggregation of fibrils

Monomeric NM (2.5 μM) were aggregated in the assembly buffer (40 mM HEPES-KOH pH 7.4, 150 mM KCl, 20 mM MgCl_2_, 1 mM DTT,10 μM ThT) without or with Hsp104 (0.025 μM), ATP (5 mM), and the ATP regeneration system (20 mM PEP, and 15 μg/ml pyruvate kinase) for 6 h or 30 h, respectively, at room temperature under stirring at 80 rpm to generate the fibrils and after that Hsp104 was added in both the reactions to a final concentration of 0.5 μM along with ATP (5 mM), and the ATP regeneration system (20 mM PEP, and 15μg/ml pyruvate kinase) and kept at room temperature under stirring at 80 rpm. A drop in the ThT fluorescence was recorded with time. Alternatively, the fibrils were also disaggregated by the ultrasonic sound (Qsonica probe sonicator) of amplitude 5 for 30 sec for several pulses, and the ThT fluorescence was recorded after each 30-sec pulse. Also, the fibrils were kept at room temperature for 24 h in the dark, static condition to record the decrease in the ThT fluorescence due to the auto-disaggregation. The percentage of disaggregation was estimated using [(Initial ThT fluorescence intensity - final ThT fluorescence intensity)/Initial ThT fluorescence intensity] × 100%.

#### Steady-state fluorescence measurements

Steady-state fluorescence measurements for tryptophan mutants of Sup35NM were performed in NM and NM-Hsp104 amyloid states using the FluoroMax-4 spectrofluorometer (Horiba Jobin Yvon, Edison, NJ). For recording the fluorescence spectra, the mutants were excited at 295 nm, where the excitation and emission slit widths were 1.75 and 6 nm, respectively. Concomitantly, steady-state anisotropy measurements were performed by setting the excitation wavelength at 295 nm and emission wavelength at 330 nm with an integration time of 2 sec and a bandpass of 2.5 nm and 10 nm, respectively. All of the above measurements were done at 24 ±1° C, and steady-state fluorescence measurements were estimated by using the parallel and perpendicular intensities taking into consideration the G-factor as shown below:^32^

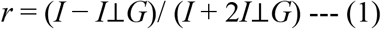

#### Atomic force microscopy (AFM)

Monomeric NM (2.5 μM) were aggregated in the assembly buffer (40 mM HEPES-KOH pH 7.4, 150 mM KCl, 20 mM MgCl_2_, 1 mM DTT) without or with Hsp104 (0.025 μM), ATP (5 mM), and the ATP regeneration system (20 mM PEP, and 15 μg/ml pyruvate kinase) and samples were aliquoted after different time points from the commencement of the reactions for imaging. For AFM imaging, the mica was freshly cleaved and washed with filtered water. Twenty μL of the sample was deposited on mica. The sample was incubated for 5 min. The mica was washed with 100 μL of filtered water twice, followed by drying under a gentle nitrogen stream. The AFM images were acquired on Innova atomic force microscopy (Bruker) using NanoDrive (v8.03) software. The images were processed and analyzed using WSxM 5.0 Develop software.^54^ The height profiles were plotted using Origin 9.65.

## Acknowledgements

We thank IISER Mohali, Department of Biotechnology (grant to S. Mukhopadhyay), Council of Scientific and Industrial Research (fellowship to P.M.), Department of Science and Technology (INSPIRE fellowship to S. Mahapatra and NanoMission grant to S. Mukhopadhyay), Ministry of Education (Centre of excellence grant to S. Mukhopadhyay) for financial support, Dr. Dominic Narang for the plasmid used in this work, Dr. Deepak Sharma (Institute of Microbial Technology, Chandigarh) for providing us with the recombinant TEV protease, and the members of the Mukhopadhyay lab for critically reading this manuscript.

## Author Contributions

S. Mahapatra, A.S., and S. Mukhopadhyay conceived the research and designed the study. S. Mahapatra and A.S. performed the experiments and analyzed the data. P.M. and H.M.S assisted in the design, experiments, and analysis. S. Mahapatra, A.S., and S. Mukhopadhyay wrote the paper. All authors discussed the results and commented on the manuscript.

