## Supporting Information for "Sub-stoichiometric Hsp104 regulates the genesis and persistence of self-replicable amyloid seeds of a yeast prion protein"

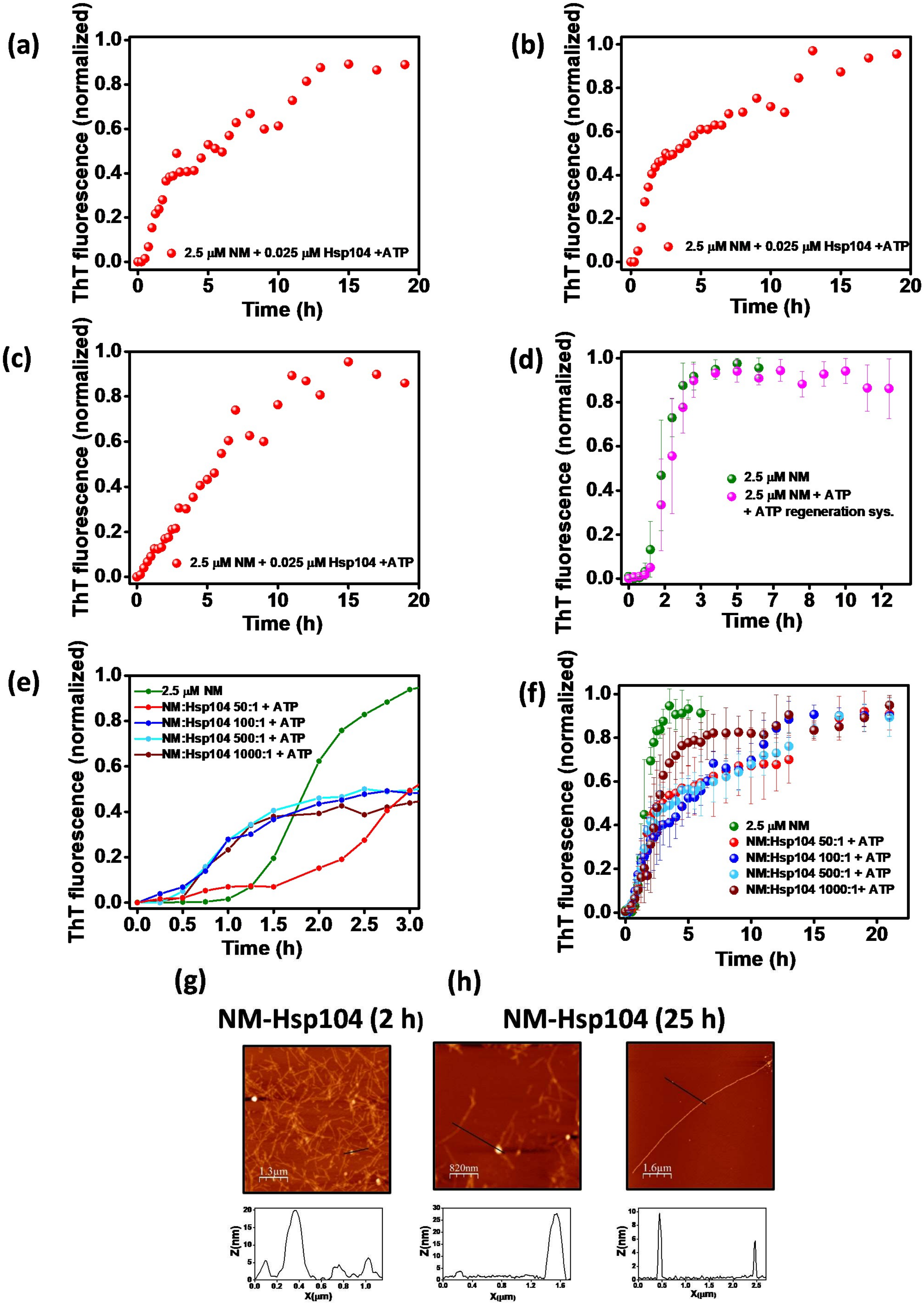
(a)
ThT fluorescence (normalized)
Time (h)
2.5 µM NM + 0.025 µM Hsp104 +ATP
(b)
ThT fluorescence (normalized)
Time (h)
2.5 µM NM + 0.025 µM Hsp104 +ATP
(c)
ThT fluorescence (normalized)
Time (h)
2.5 µM NM + 0.025 µM Hsp104 +ATP
(d)
ThT fluorescence (normalized)
Time (h)
2.5 µM NM
2.5 µM NM + ATP
+ ATP regeneration sys.
(e)
ThT fluorescence (normalized)
Time (h)
2.5 µM NM
NM:Hsp104 50:1 + ATP
NM:Hsp104 100:1 + ATP
NM:Hsp104 500:1 + ATP
NM:Hsp104 1000:1 + ATP
(f)
ThT fluorescence (normalized)
Time (h)
2.5 µM NM
NM:Hsp104 50:1 + ATP
NM:Hsp104 100:1 + ATP
NM:Hsp104 500:1 + ATP
NM:Hsp104 1000:1+ ATP
(g)
(h)
NM-Hsp104 (2 h)
NM-Hsp104 (25 h)
1.3µm
820nm
1.6µm
Z(nm)
X(µm)

**Figure S1.** (a-c) Representative normalized rotated (80 rpm) thioflavin-T (ThT) fluorescence kinetics of NM (2.5 μM) with Hsp104 (0.025 μM), plus ATP (5 mM) during amyloid formation at room temperature. (d) Normalized ThT fluorescence kinetics of rotated (80 rpm) NM (2.5 μM) aggregation without or with ATP (5 mM) and ATP regeneration system [Phosphoenolpyruvate (20 mM) and pyruvate kinase (15 μg/ml)]. (e,f) Normalized ThT fluorescence kinetics of NM (2.5 μM) without or with Hsp104 and ATP (5 mM) during amyloid formation at room temperature and 80 rpm showing the (e) first 3 h (representative kinetics) and (f) 20 h of aggregation. The standard deviations were estimated from three independent experiments (g,h) AFM images of the (g) NM oligomers and protofibrils with the height ~ 20 nm and 7 nm, respectively after 2 h and (h) NM oligomers, protofibrils, and fibrils formed after 25 h of aggregation with the height ~ 25 nm,~5 nm, and ~10 nm, respectively in the presence of Hsp104 (0.025 μM) and ATP.

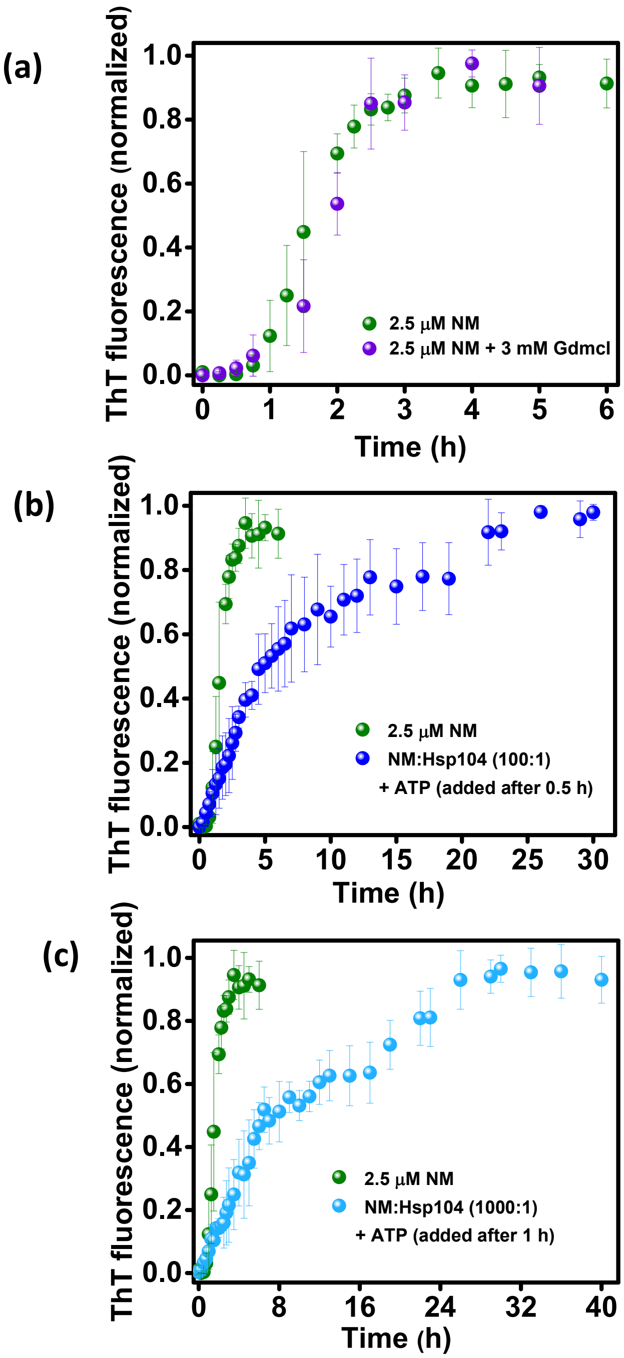


**Figure S2.** (a) Normalized ThT fluorescence kinetics of rotated (80 rpm) NM (2.5 μM) aggregation without or with GdmCl (3 mM) during amyloid formation. (b) Normalized ThT fluorescence kinetics of rotated (80 rpm) NM (2.5 μM) aggregation without or with Hsp104 (0.025 μM), plus ATP, both introduced after 0.5 h from the commencement of the reaction. (c) Normalized ThT fluorescence kinetics of rotated (80 rpm) NM (2.5 μM) aggregation without or with Hsp104 (0.0025 μM), plus ATP, both introduced after 1 h from the commencement of the reaction. (All the standard deviations were calculated from three independent experiments).

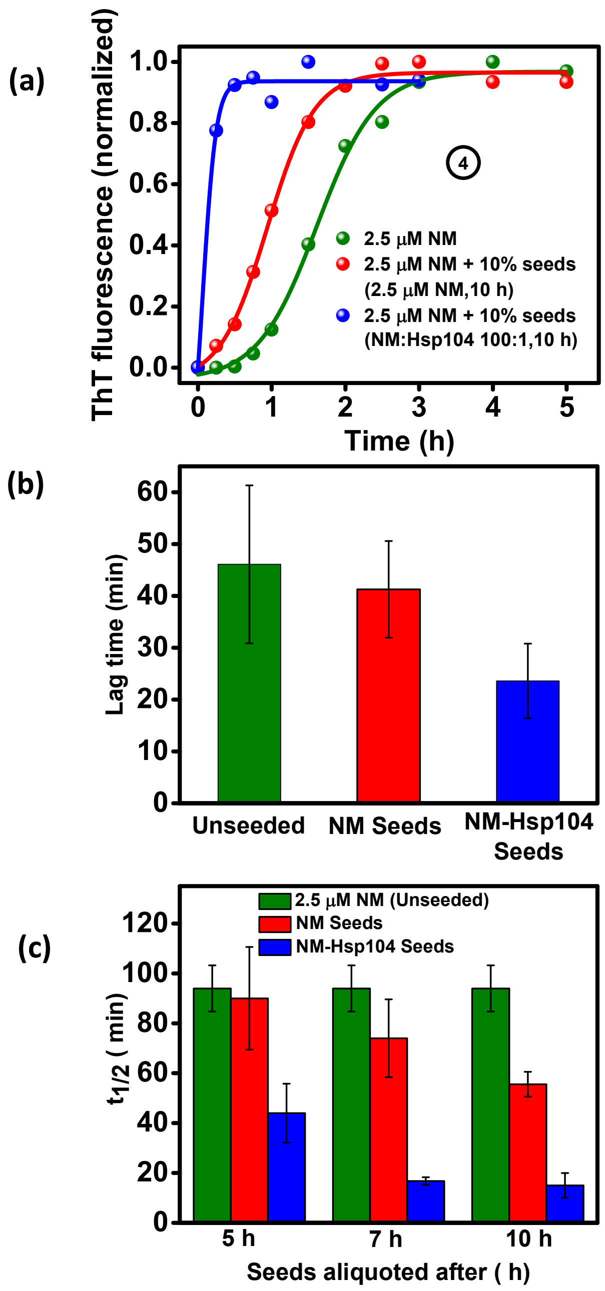


**Figure S3.** (a) Representative normalized ThT fluorescence kinetics of rotated (80 rpm) NM (2.5 μM) aggregation without or with 10% (w/w) seeds of NM-Hsp104 or NM aggregation that aliquoted after 10 h from the commencement of the aggregation reaction. (b) From figure 3b, lag times of the unseeded aggregation reactions and aggregation reactions seeded with amyloids from NM or NM-Hsp104 aggregation reaction were recovered from the fitted sigmoidal plots. (c) From figure 3c,d and S3a, $t_{1/2}$ of the unseeded or aggregation reactions seeded with amyloids from NM and NM-Hsp104 aggregation reactions formed after 5 h, 7 h, and 10 h from the commencement of reactions. (All the standard deviations were estimated from three independent experiments.)

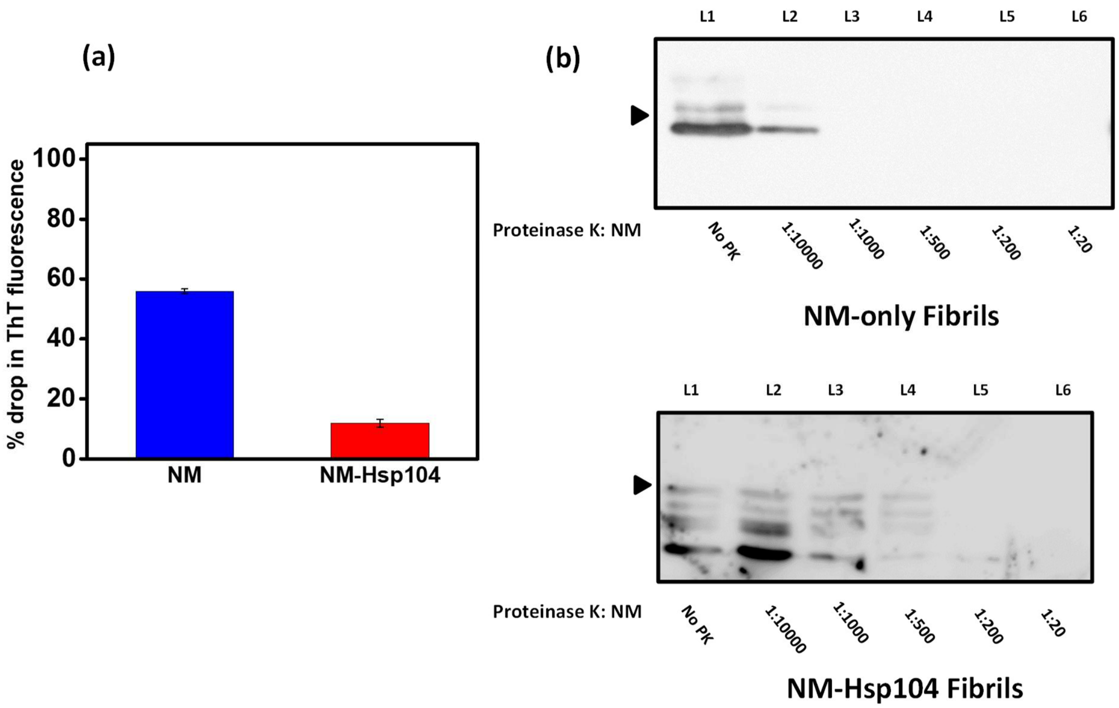


**Figure S4.** (a) Auto-disaggregation of NM and NM-Hsp104 fibrils by keeping the fibrils at room temperature for 24 h under the unagitated condition and the drop in the ThT fluorescence was estimated. (b) The concentrated fibrils formed from monomeric NM (2.5 µM) without or with Hsp104 (0.025 µM), plus ATP were incubated at 37 °C for 30 min with multiple concentrations of proteinase K followed by the western blot analysis with the anti-His antibody. The bands corresponding to the NM monomers are marked with black triangles.

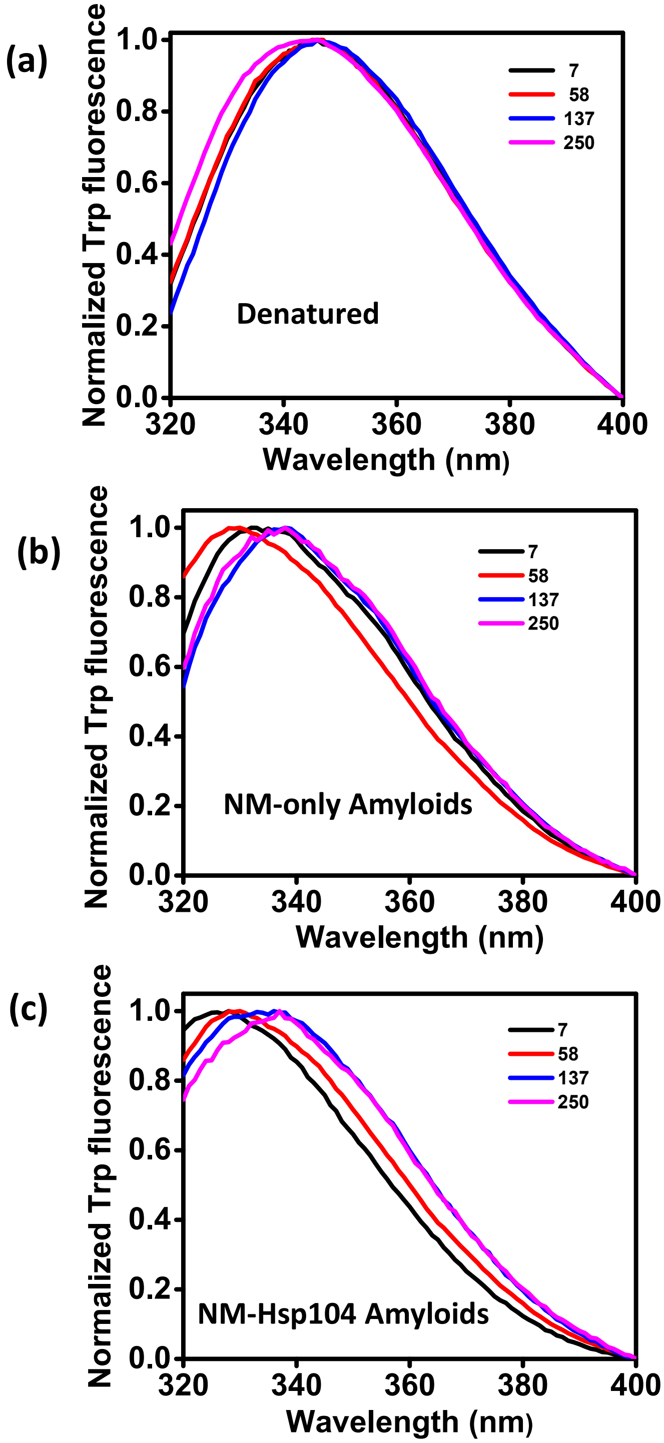


**Figure S5.** Normalized Trp fluorescence spectra of different residues in (a) denatured (b) NM-only amyloid (c) NM-Hsp104 amyloid states.
